# Infectious touching: Has COVID-19 changed our perceptions of social touch? A neural and behavioral study

**DOI:** 10.1101/2023.09.12.557330

**Authors:** Dana Zoabi, Elinor Abado, Simone Shamay-Tsoory, Leehe Peled-Avron

**Affiliations:** School of Psychological Sciences, University of Haifa, Haifa, Israel; The Integrated Brain and Behavior Research Center (IBBRC), University of Haifa, Haifa, Israel; Psychology Department and Gonda Brain Research Center, Bar Ilan University, Israel

## Abstract

Social touch is essential for reducing stress, improving mood and fostering a sense of social connectedness. Stimuli related to social touch are generally perceived as positive. Nevertheless, the social restrictions imposed by the COVID-19 pandemic may have changed the way human beings perceive and react to social touch. Indeed, the social distancing imposed by the pandemic may have had long term effects on human perceptions of social touch. In the current study, we examined how perceptions of interpersonal touch in social interactions were affected by the COVID-19 pandemic. Specifically, we compared behavioral and neural responses to observed social touch between two groups: pre- and post-COVID-19. Participants in both groups rated the pleasantness of photos of social touch between humans, nonsocial touch between inanimate objects or non-touch photos of either two humans or inanimate objects. We hypothesized that social touch in the post-COVID-19 group would induce hypervigilance due to the risk of infection. In line with our predictions, we found behavioral changes in perceptions of social touch among participants in this group, who rated photos with touch as less pleasant than did participants in the pre-COVID-19 group. Participants in the post-COVID-19 group also rated photos with humans as less pleasant than did participants in the pre-COVID-19 group. Additionally, EEG analysis revealed neural changes in the ERP components associated with hypervigilance: P1 and LPP. Contrary to pre-COVID-19 measures showing more positive P1 amplitudes for touch than for non-touch photos, after COVID-19 no differences in P1 amplitudes were found between touch and non-touch photos. Furthermore, after COVID-19 the P1 amplitudes for human and inanimate photos in the touch condition were similar, a pattern that did not emerge prior to COVID-19. These findings suggest that COVID-19 had a surprising impact on human perceptions of social touch, such that observing nonsocial touch evoked more positive emotions than observing human touch. Further, these findings may reflect shifts in attention or changes in the salience of touch-related information due to the altered circumstances brought about by the pandemic. Overall, our results indicate that COVID-19 has modified human perceptions of social touch, providing evidence that the pandemic has affected individuals’ perceptual and evaluative processes and highlighting the importance of considering social and environmental factors in understanding subjective experiences.

## Background

Social touch is defined as physical contact between human beings. It includes gestures such as hugging, holding hands, stroking or patting someone on the back and can occur in various social interactions (Cascio et al., 2019). Touch is a fundamental human need and plays a crucial role in social interactions and communication. It can convey a wide range of emotions (McIntyre et al., 2022) and has been linked to physical and mental health benefits such as reduced stress and improved mood (Cascio et al., 2019; Hertenstein et al., 2009; Hertenstein et al., 2007) and an enhanced sense of social connectedness to others (Eckstein et al., 2020). Given the magnitude of the effects of social touch, scholars have claimed that the mere observation of touch produces positive effects. For example, Schirmer et al., (2015) reported that observing social touch is associated with positive emotions and that characters in photos appear more positive and likeable when they touch each other than when they do not. In line with the pleasant effects of observed touch, Peled-Avron et al. (2016) examined whether interindividual variations in empathy levels affected participants’ reactions to observing touch-based social interactions. Their findings indicated that participants rated images of human social touch as evoking more pleasant emotions than images depicting nonsocial touch (between inanimate objects) or no touch, regardless of their empathy levels (high or low).

In an attempt to examine the neural processing of observed touch, scholars have studied event-related potentials (ERPs) (reviewed in Peled-Avron & Woolley, 2022). ERPs are brain-generated electrical potentials that are associated with internal or external events (e.g., stimuli, responses, decisions; Bressler & Ding, 2006). *P1* is an early ERP component that occurs approximately 110ms after stimulus onset, is measured at parietal sites and reflects attention allocation (Doesburg et al., 2008; Wijers et al., 1997). During tasks involving stimulus discrimination, the amplitude of the P1 component increases when the participant concentrates on the stimulus. This heightened amplitude signifies a reduction in processing at unattended locations and minimized interference between attended and unattended information. Thus, the P1 component serves as a direct indicator of attentional resource allocation and effort, making it a neural marker for selective spatial attention (Heinze et al., 1994). P1 was found to have shorter latencies in response to social touch than to non-social touch stimuli, suggesting that participants allocate more attentional resources to social touch (Peled-Avron & Shamay-Tsoory, 2017). This result implies that observed touch draws people’s attention away from other non-social stimuli.

Another ERP component that researchers suggested to be activated during observed touch is the *late positive potential (LPP)* component (Peled-Avron & Shamay-Tsoory, 2017). LPP has a positive amplitude that peaks 400-600ms post stimulus onset (Peyk et al., 2006; Schupp et al., 2000, 2003). LPP is typically measured at parietal electrodes and is higher in response to emotional stimuli than neutral stimuli (Hajcak et al., 2010; Schupp et al., 2000). Unlike the P1 component, *LPP* is associated with the processing of higher and more complex cognitive and emotional information, such as affective images (Olofsson et al., 2008), human voices (Schirmer & Gunter, 2017), facial expressions (Hartigan & Richards, 2016) and felt touch (Ackerley et al., 2013). This component is thought to measure motivated attention toward stimuli because it persists throughout the duration of stimulus presentation. Increased LPP amplitudes have been demonstrated in response to observed touch (Adler & Gillmeister, 2019; Peled-Avron & Shamay-Tsoory, 2017; Schirmer & McGlone, 2019), possibly reflecting enhanced social-emotional processing in response to observed touch.

Peled-Avron and Shamay-Tsoory (2017) investigated the neural correlates of interpersonal touch perception in individuals with high or low autistic traits by focusing on the P1 and LPP components of the ERP. In line with previous studies showing that individuals with autism spectrum disorder (ASD) have a strong aversion to interpersonal touch, these researchers found that higher autistic traits, as opposed to low autistic traits, were associated with higher LPP amplitudes, which serve as a marker of anxiety bias, and with shorter P1 latencies, which serve as a marker for detection and allocation of attention to social touch. The early ERP component (P1) appears to reflect enhanced allocation of attentional resources to stimuli, whereas the later ERP components (LPP) reflect emotional and social reactions to observed touch (Peled-Avron & Woolley, 2022).

Many social factors can affect human perception of social touch (Sailer & Leknes, 2022). For instance, the recent COVID-19 pandemic was found to have a major impact on how people perceive and interpret social touch (Von Mohr et al., 2021). COVID-19 is a severe respiratory disease that first appeared in 2019 and spread rapidly across the globe. It caused thousands of deaths, thus prompting the World Health Organization to declare it as a pandemic in March 2020. The COVID-19 pandemic brought about significant changes in social behavior as people had to adapt to new norms of social restrictions and keep their distance from one another to prevent the spread of the disease (Abbasi, 2020). All ordinary social behaviors, such as meeting with friends and family members, hugging and shaking hands suddenly became unacceptable and less prevalent (Bavel et al., 2020). Restrictions on these behaviors increased levels of social isolation and loneliness (Horigian et al., 2021). Quarantine measures, physical separation and the closure of public spaces all reduced opportunities for social interaction and support. Indeed, touch—one of the factors shown to diminish loneliness and increase social connectedness (Heatley Tejada et al., 2020)—was less available during the pandemic.

In the current study, we sought to examine whether the pandemic has influenced people’s perceptions of social touch. Hence, the aim of the present study was to investigate behavioral and neural changes in perception of interpersonal touch brought about by COVID-19. What we know about perceived interpersonal touch following COVID-19 is limited. The assumption was that COVID-19 would have an impact on people’s attitudes toward social touch and perceived social touch. Yet the question remains as to whether social restrictions may have increased or decreased the pleasantness associated with social touch. While behavioral reactions to social touch were studied to some extent during and after the pandemic, the neural underpinnings of this behavior have not been studied. Therefore, this study aims to characterize the behavioral *and* neural changes in perception of interpersonal touch brought on by COVID-19 by comparing data collected before the pandemic (for behavioral data see Peled-Avron et al., (2016) and Peled-Avron & Shamay-Tsoory (2017) for neural data) to data collected afterwards. This comparison can advance our knowledge regarding the effects of COVID-19 on perceived social touch so as to enable us to better understand the impact of the pandemic on people’s social well-being and inform efforts to mitigate the negative effects of isolation and loneliness.

In view of the reduction in social touch during the pandemic and the negative reactions to social interactions shown in previous studies, we hypothesized that there would be differences between pre- and post-COVID-19 responses to observed touch, such that participants in the post-COVID-19 group would rate photos depicting social touch as less pleasant than the previous results reported by Peled-Avron et al., (2016). To our knowledge, only Kühne et al. (2022) examined observed touch between two individuals (and not a large gathering) following COVID-19, showing that participants perceived non-touch photos as more pleasant than touch photos. Based on these findings, we predict that our data will reflect such a pattern. We further hypothesize that this change will also be reflected in electrophysiological activity. Compared to pre-COVID-19, we expected that post-COVID-19 LPP amplitudes would be lower in response to social touch, reflecting heightened emotional reactions or heightened anxiety elicited by social touch. We also hypothesized that compared to pre-COVID-19, P1 amplitude would be lower, indicating reduced attentional allocation or sensitivity to social touch stimuli. This reduction may reflect decreased significance of social touch information or less interest in processing such information following COVID-19. We also hypothesized that P1 latency in response to social touch in the post-COVID-19 group would be greater than in the pre-COVID-19 group. Such an increase in latency would suggest delayed or slower initial processing of social touch information, possibly reflecting a lower inclination or heightened caution when engaging in social touch interactions following COVID-19.

## Methods

### Participants

Forty-three participants (25 female) were recruited from the University of Haifa, Israel. They received course credit or payment in exchange for participating in the experiment. The participants’ ages ranged from 18 to 42 (*M*= 26.15, *SD*± 5.813). Five participants were excluded from the neural data analyses due to failure to clear substantial noise from their data, leaving 38 participants for the neural data analysis. All participants were right-handed and reported normal or corrected-to-normal vision. A screening interview confirmed that all participants had no history of psychiatric or neurological disorders. Participants gave their signed informed consent to participate in the study. This study used the same participant selection criteria as described in Peled-Avron et al. (2016).

### Stimuli Task and Design

The computerized task used in this experiment is similar to the one described by Peled-Avron et al. (2016). The task was administered to participants who were seated about 60 cm from a 21" CRT monitor. E-Prime 2.2 Psychological Software was used for stimulus presentation and experimental control. To account for any potential low-level visual changes across the stimulus categories, participants were shown monochromatic images that were 15 cm × 10 cm in size and had fixed luminance (Johannes et al., 1995).

A total of 45 photos were shown, divided into four conditions—humans or objects either touching or not touching—forming a 2×2 design of object (human, inanimate) × touch (with, without). The human touch images depicted a hug, a handshake, a tap on the shoulder or friendly handholding (Figure 1a). The inanimate contact pictures depicted two common objects without any commercial logos (e.g., cutlery, clothing items) touching one another. In the non-touch condition, humans or objects were shown without touching. This combination of touch and object type yielded a 2 × 2 factorial design.

**Figure 1.**
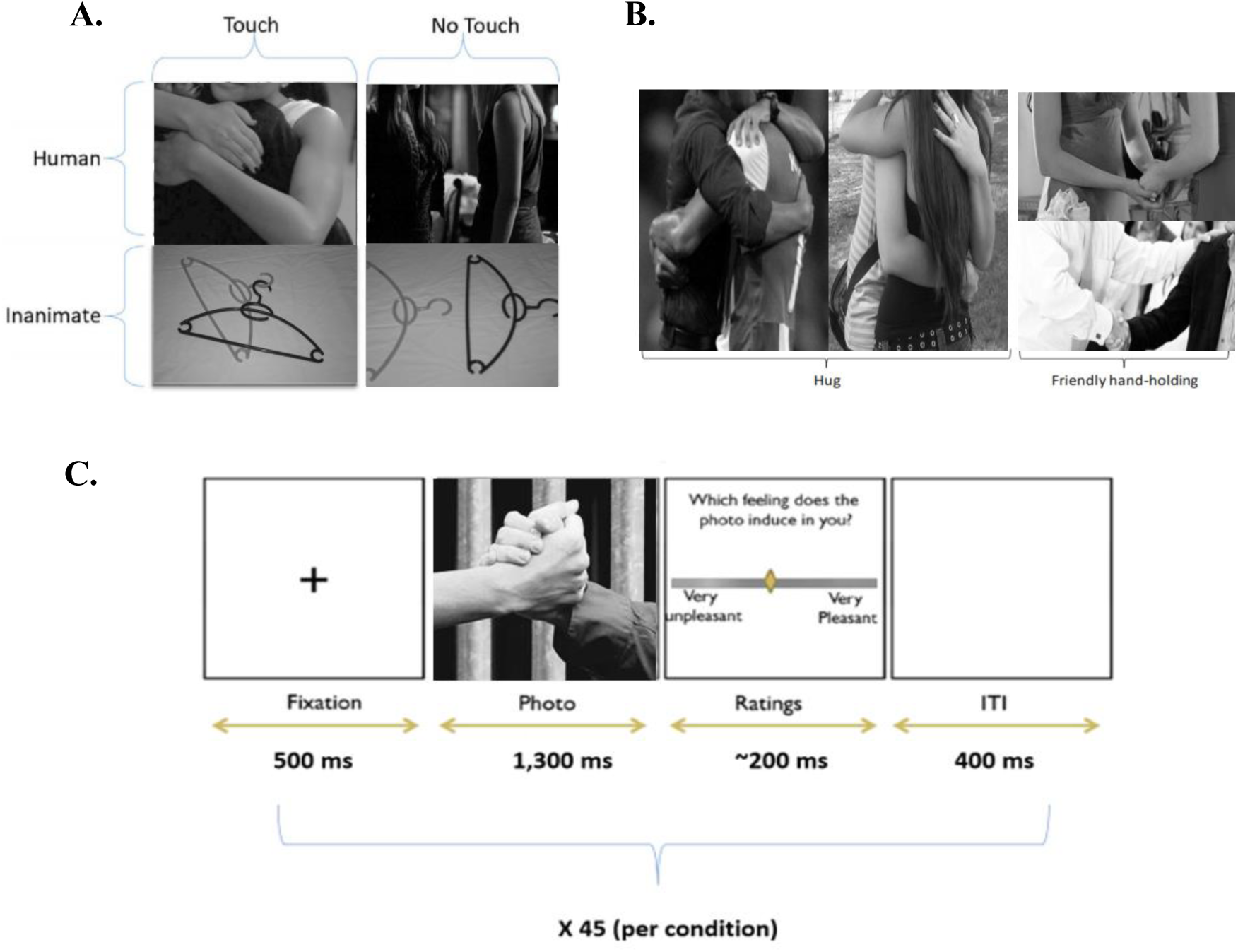
Examples of the task stimuli and design. (**A**) Sample stimuli images depicting a natural social interaction in a public setting between two women that did or did not involve touch and control images depicting two inanimate objects either touching or not. (**B**) Sample images depicting social interactions involving touch between two males and two females. (**C**) Trial scheme in the computerized paradigm: A fixation cross followed by the photo and rating screen, with an interstimulus interval between each trial. Each condition contains 45 photos counterbalanced between four blocks. ITI=intertrial interval.

Participants were instructed to rate the extent of pleasantness or unpleasantness evoked by each photo. We employed a bipolar valence visual analog scale (VAS) to quantify participants’ subjective responses. Subsequently, the VAS ratings were converted into numerical values offline, ranging from 0 (representing unpleasant feelings) to 100 (representing pleasant feelings). In accordance with the IAPS protocol (Lang et al., 1998), participants were instructed as follows: "At one extreme of the scale, you feel completely unpleasant, unhappy, annoyed, dissatisfied, melancholic or despairing. On the other extreme, you experience complete satisfaction, joy, happiness, contentment, or hope”. Further, participants were informed that a score at the scale’s middle point represented a completely neutral state that was neither pleasant/happy nor unpleasant/sad.

As described in Peled-Avron et al. (2016), male participants were shown images depicting social touching between men, whereas female participants were shown images depicting social touching between women (see Figure 1b). Moreover, all the portrayed social interactions took place between same-sex friends. The human condition photographs depicted the neck and down with no faces shown in order to keep the stimuli simple and reduce any potential confounding effects. The stimuli were presented in four blocks of 45 trials each, for a total of 180 trials. Every trial started with a fixation cross shown for 500 ms, followed by a picture for 1,300 ms, and an inter-trial period of a blank screen for 400 ms. Each participant was shown a different set of photos in each of the four blocks, in random order (see Figure 1c).

### EEG Data acquisition

During the behavioral task, EEG was recorded from 32 scalp sites using active, gel-based Ag/AgCl electrodes mounted in an elastic cap (g.Nautilus PRO, g.Tec medical engineering GmbH, Austria, https://www.gtec.at/product/gnautilus-pro/) using an extended 10–20 system. The EEG signals were digitally amplified and sampled at 500 Hz. Data were wirelessly transmitted using 2.4 GHz band. Impedances were maintained below 5 kΩ.

### Behavior analysis

To test participants’ emotional response to the four conditions, we conducted a repeated-measures analysis of variance (ANOVA), with *group* as the between-subjects factor (pre-COVID-19 and post-COVID-19) and *touch* (touch, no touch) and *object* (human, inanimate) as within-subject factors. The emotional ratings of the photos served as the dependent variable. Behavioral data were analyzed using IBM SPSS Statistics (version 27.0.1).

### EEG Data Processing

Data were analyzed using EEGLab (version 2021.0; Delorme & Makeig, 2004) and ERPLab plugin (Lopez-Calderon et al., 2014) running on MATLAB (MathWorks; version R2021b) routines. Raw EEG data were re-referenced offline to the digital average of the 32 EEG electrodes. EEG deflections resulting from eye blinks were corrected using independent component analysis (ICA). Any remaining artifacts that exceeded ±100 µV in amplitude were rejected.

### ERP Analysis

ERPs were determined by averaging the 1300-ms segmented trials separately in each stimulus condition (human touch, human non-touch, object touch, and object non-touch). The averaged waveforms were smoothed by applying a 30-Hz low-pass filter and were baseline corrected to the signal recorded 200ms before stimulus onset. Based on LPP literature discussing visual processing of the affective content in photos (Peyk et al., 2006; Schupp et al., 2000, 2003), LPPs were analyzed for the vertex electrodes (Pz, Cz, and Fz; Schupp et al., 2003). LPP amplitudes were scored as the mean amplitude in the time interval from 400 to 600ms following stimulus onset. The statistical analysis of the P1 component was limited to the parietal sites P7 and P8, which are commonly used for P1 analyses (Doesburg et al., 2008; Wijers et al., 1997). The peak of the P1 component was defined for each participant as the largest positive peak amplitude between 100 and 150ms. Visual inspection ensured that these values represented actual peaks rather than epoch end-points. LPP, P1 amplitudes and P1 latency were compared before and after COVID-19. Interactions with type of photo were assessed using repeated measures of variance (ANOVA), with *group* as the between-subjects factor (pre-COVID-19 and post-COVID-19) and *touch* (touch, no touch) and *object* (human, inanimate) as within-subject factors. The dependent variable was the LPP amplitude, P1 amplitude or P1 latency.

## Results

### Behavior

To examine the hypothesis that perception of social touch was affected by COVID-19, we conducted a repeated-measures analysis of the subjective ratings, with *touch* (touch, no-touch) and *object* (human, inanimate) as within-subject factors and *COVID-19* (pre-COVID-19, post-COVID-19) as a between-subjects factor.

The subjective ratings analysis revealed a significant three-way interaction between touch, object, and COVID-19 (*F* (1, 106) =33.399, *P*<.001, partial *η^2^*=.24; see Figure 2). To understand this interaction, we examined the pre- and post-COVID-19 conditions separately. In the pre-COVID-19 condition, an interaction emerged between touch and object *(F* (1,64) =77.91, *P*<.001, partial *η^2^* =.549); in the touch condition, the subjective ratings of human photos were higher than those of inanimate photos (*P*<.001), while no differences were found in the non-touch condition (*P*=.205). Furthermore, a main effect emerged for touch (*F* (1,64) =46.827, *P*<.001, partial *η^2^*= .423) as well as for object (*F* (1,64) = 99.048, *P*<.001, partial *η^2^*= .607). These findings indicate that prior to COVID-19 participants showed a preference for photos with touch, perceiving them as more pleasant than non-touch photos *(P*<.001). Additionally, photos featuring humans were rated as more pleasant than those showing inanimate objects (*P*<.001). In the post-COVID-19 condition as well, the interaction between touch and object was significant (*F* (1,42) = 4.004, *P*=.052, partial *η^2^*= .087): For the touch condition, no differences emerged between photos with human touch and photos with inanimate touch (*P*=.466), whereas in the non-touch condition, participants assigned higher subjective ratings to human photos (*P*=.012).

**Figure 2.**
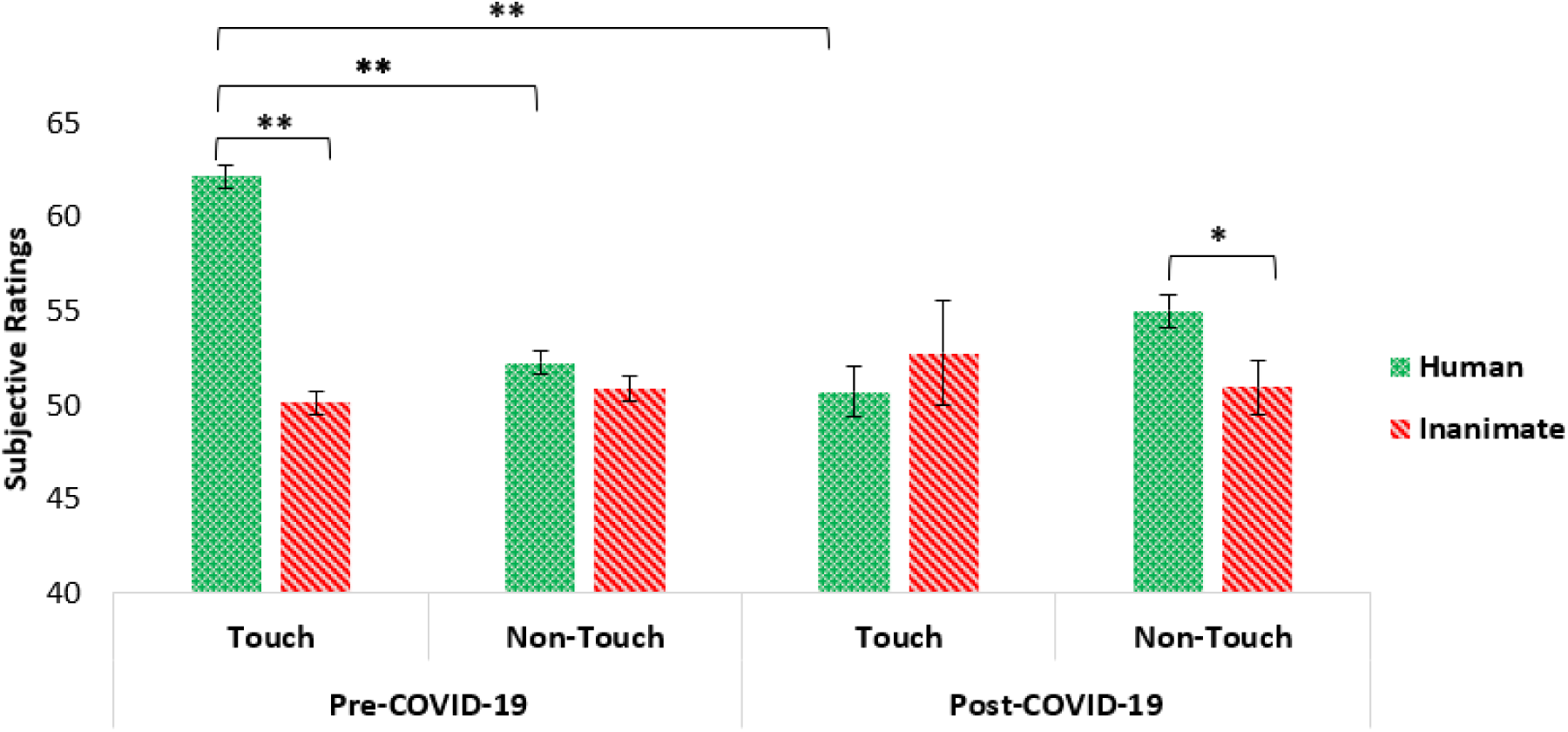
Subjective behavioral ratings for the pre- and post-COVID-19 groups. Participants in the post-COVID-19 group rated human touch photos as inducing less pleasant emotions than did participants in the pre-COVID-19 group. In the pre-COVID-19 group, subjective ratings of human touch photos were higher than ratings both of inanimate touch photos and of non-touch photos. In the post-COVID-19 group, however, participants rated human non-touch as more pleasant than inanimate non-touch. Main effects were found for the touch condition and the object condition. *Error bars represent standard error of the mean. *P<.05 **P<.001*

In accordance with our hypothesis, the analysis revealed a significant interaction between touch and COVID-19 (*F* (1,106) = 11.192, *p* = .001, partial *η^2^*=.096), indicating that participants in the post-COVID-19 group rated photos depicting touch as less pleasant than did participants in the pre-COVID-19 group (*F* (1,42) =8.385, *P*=.005, partial *η^2^*=.073). In the non-touch conditions, no differences emerged between the pre- and post-COVID-19 groups (*F* (1,64) =2.746, *P*=.100, partial *η^2^*=.025).

Moreover, the analysis revealed a significant interaction between object and COVID-19 (*F* (1,106) = 12.771, *p* = .001, partial *η^2^*=.108). Thus, participants in the pre-COVID-19 group rated photos with humans as more pleasant than did participants in the post-COVID-19 group (*F* (1,42) =10.452, *P*=.002, partial *η^2^*=.09). For the inanimate condition, no differences were found between the pre- and post-COVID-19 groups (*F* (1,64) =1.816, *P*=.181, partial *η^2^*=.017). In addition, we found a significant main effect for the touch condition (*F* (1, 106) = 3.689, *p* =.057, partial *η^2^*=.032), such that participants rated photos containing touch as inducing more pleasant emotions than non-touch photos. A significant main effect for object emerged as well (*F* (1, 106) =23.488, *p* <.001, partial *η^2^*=.181), such that photos containing humans were rated as inducing more pleasant emotions than photos containing inanimate objects.

### P1 amplitude

To test whether COVID-19 affected P1 amplitudes while participants viewed pictures depicting touch, we conducted a repeated-measures analysis, with *touch* (touch, no-touch) and *object* (human, inanimate) as within-subject factors and *COVID-19* (pre-COVID-19, post-COVID-19) as a between-subjects factor. The analysis revealed a significant three-way interaction between touch, object, and COVID-19 (*F* (1,90) =5.972, *p* =.016, partial *η^2^* =.062; see Figures 3, 4). To gain a better understanding of the interaction, we examined the COVID-19 conditions separately. In the pre-COVID-19 condition, no interaction was found between touch and object *(F* (1,53) =2.24, *P*= .14, partial *η^2^*= .041), while a main effect emerged for touch and for object *(F* (1,53) =42.423, *P*<.001, partial *η^2^*= .445; *F* (1,53) = 20.866, *P*<.001, partial *η^2^*= .282; respectively). Furthermore, in the pre-COVID-19 condition, P1 amplitudes were more positive when participants viewed photos depicting touch compared to photos with no touch and when they viewed human photos compared to inanimate object photos.

**Figure 3.**
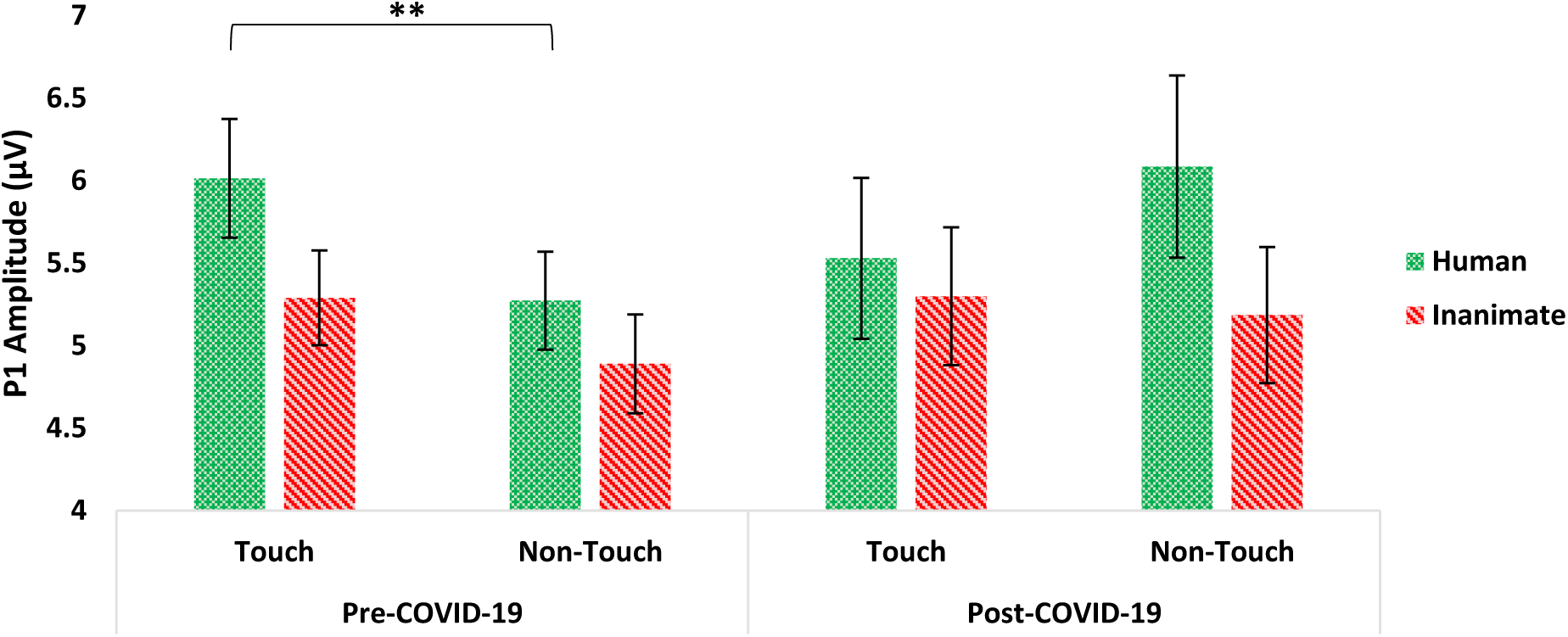
P1 amplitudes for the pre- and post-COVID-19 groups (averaged between P7 and P8 electrodes, 100–150ms post-stimulus). In the pre-COVID-19 group, the P1 amplitudes showed a more positive response when viewing photos with touch compared to photos without touch. Yet after the pandemic no significant differences emerged between touch photos and non-touch photos. Main effects for both touch and objects were found. *Error bars represent standard error of the mean **P<.001*

**Figure 4.**
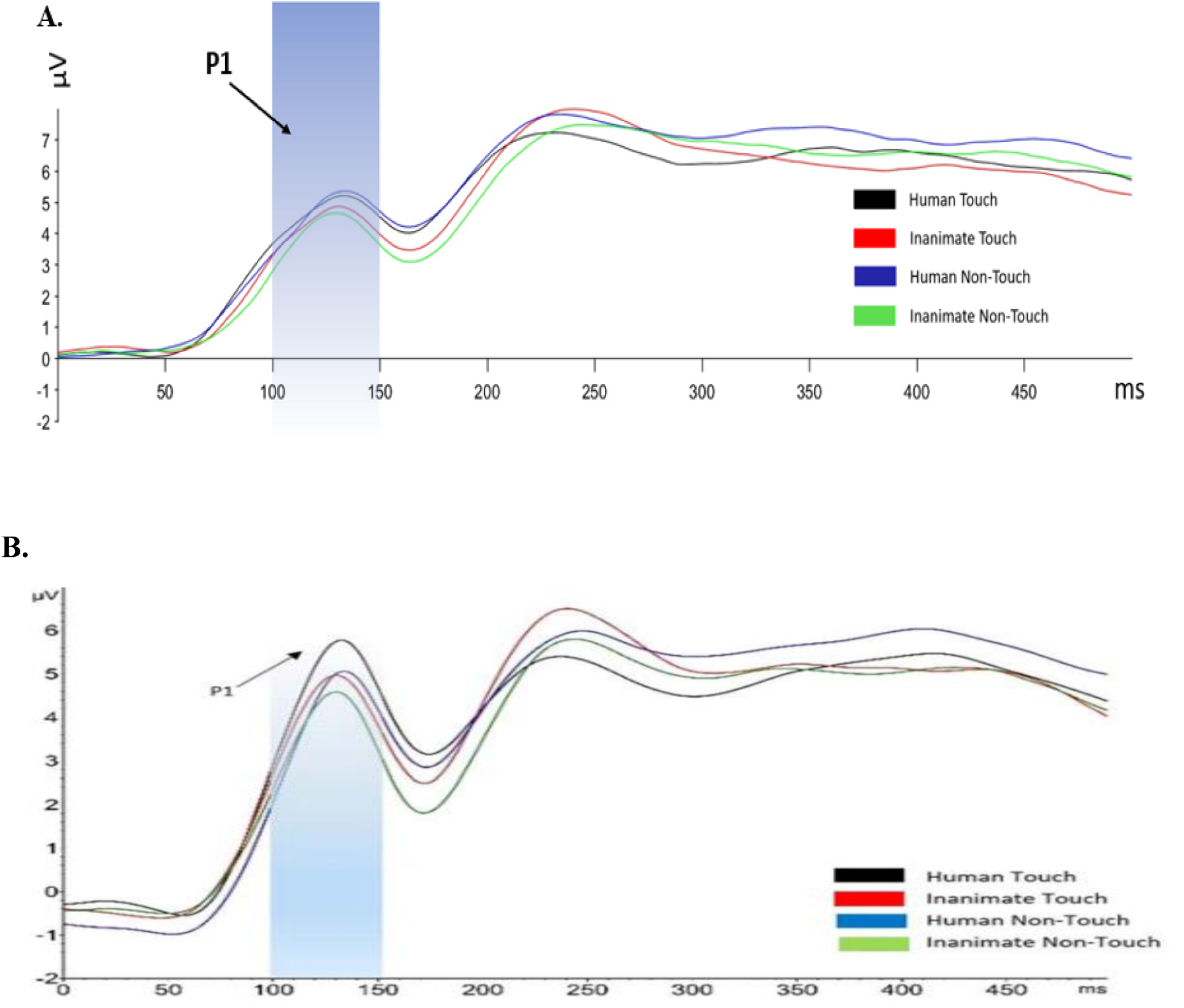
Onset-synchronized grand-average ERP P1 component measured under the indicated experimental conditions. The P1 component was averaged between P7 and P8 electrodes. The blue shaded areas represent the time windows for P1 (approximately 130ms post-stimulus). **(A)** P1 amplitudes for the post-COVID-19 group; **(B)** P1 amplitudes for the pre-COVID-19 group (adapted from Peled-Avron et al., 2016).

In the post-COVID-19 condition, no interaction was found between touch and object (*F* (1,37) = 3.236, *P*=.08, partial *η^2^*= .08). As shown in Figure 5, a main effect emerged for object (*F* (1,37) = 8.106, *P*= .007, partial *η^2^*= .18), such that P1 amplitudes were more positive when participants viewed human photos than when they viewed inanimate object photos.

**Figure 5.**
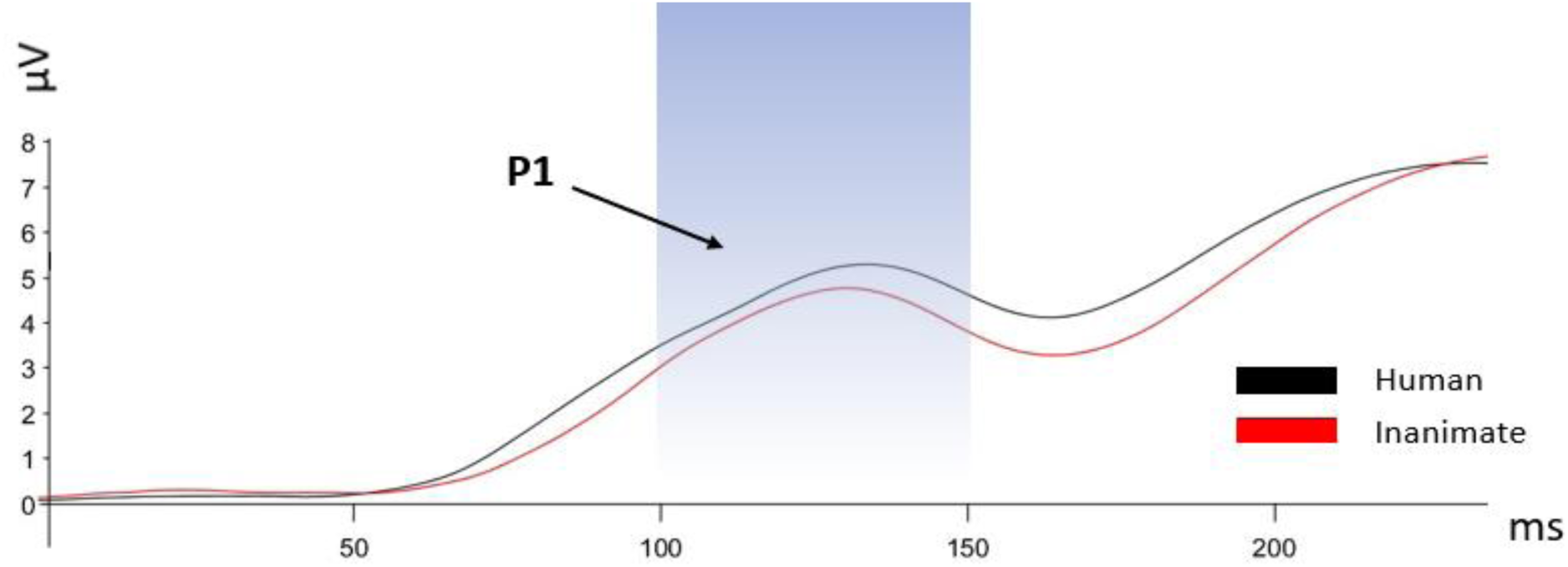
Onset-synchronized grand-average ERP P1 waveforms post-COVID-19 for the human (black) and inanimate (red) conditions measured at electrodes P7 and P8. The blue shaded area indicates the time window for P1 (approximately 130ms post-stimulus). The P1 amplitude was significantly shorter for the inanimate objects condition *(P <.05)*.

No significant interaction emerged between object and COVID-19 (*F* (1,90) = 0.003, *p* =.95, partial *η^2^* =.00). Nevertheless, a significant interaction between touch and COVID-19 emerged (*F*(1,90) = 20.85, *p* <.001, partial *η^2^* =.188); Specifically, in the pre-COVID-19 group the P1 amplitudes were more positive toward touch photos than when they viewed non-touch photos (*F* (1,53) = 26.238, *p* <.001, partial *η^2^* =.226), while in the post-COVID-19 group no differences were found between touch and non-touch photos (*F*(1,37) = 2.766, *p* = .1, partial *η^2^* =.03).

In addition, a significant main effect emerged for touch (*F* (1, 90) = 4.072, *p* =.047, partial *η^2^*=.046) and for object (*F* (1, 90) = 25.787, *p* <.001, partial *η^2^*=. 222). Specifically, P1 amplitudes were more positive when participants viewed touch photos than when they viewed non-touch photos, while photos containing humans induced more positive P1 amplitudes than photos with inanimate objects.

### P1 latency

We conducted a repeated-measures analysis to test whether COVID-19 affected P1 latency, with *touch* (touch, no-touch) and *object* (human, inanimate) as within-subject factors and *COVID-19* (pre-COVID-19, post-COVID-19) as a between-subjects factor. No three-way interaction was found between touch, object, and COVID-19 (*F* (1,90) =1.458, *p* =.23, partial *η^2^* =.016). No other interactions or main effects were found (*p*>.05).

### LPP

To examine the hypothesis that the average LPP amplitude was affected by COVID-19 we conducted a repeated-measures analysis, with *touch* (touch, no-touch) and *object* (human, inanimate) as within-subject factors and *COVID-19* (pre-COVID-19, post-COVID-19) as a between-subjects factor. LPP analysis revealed a significant three-way interaction between touch, object, and COVID-19 (*F* (1,90) =8.87, *p* =.004, partial *η^2^* =.09; see Figures 6, 7). COVID-19 groups were examined separately to gain a better understanding of this interaction. In both the pre- and post-COVID-19 groups, a significant interaction emerged between touch and object *(F* (1,53) =18.856, *P*<.001, partial *η^2^*= .262; *F* (1,37) = 29.29, *P*<.001, partial *η^2^*= .442, respectively). In the touch condition, higher LPP amplitudes were found when participants observed human touch photos compared to inanimate touch photos (*P*=.013; *P*<.001). Conversely, in the non-touch condition, the LPP amplitudes were more positive toward inanimate photos than when they observed human photos (*P*=.05; *P*=.039) in both groups (pre- and post-COVID-19, respectively).

**Figure 6.**
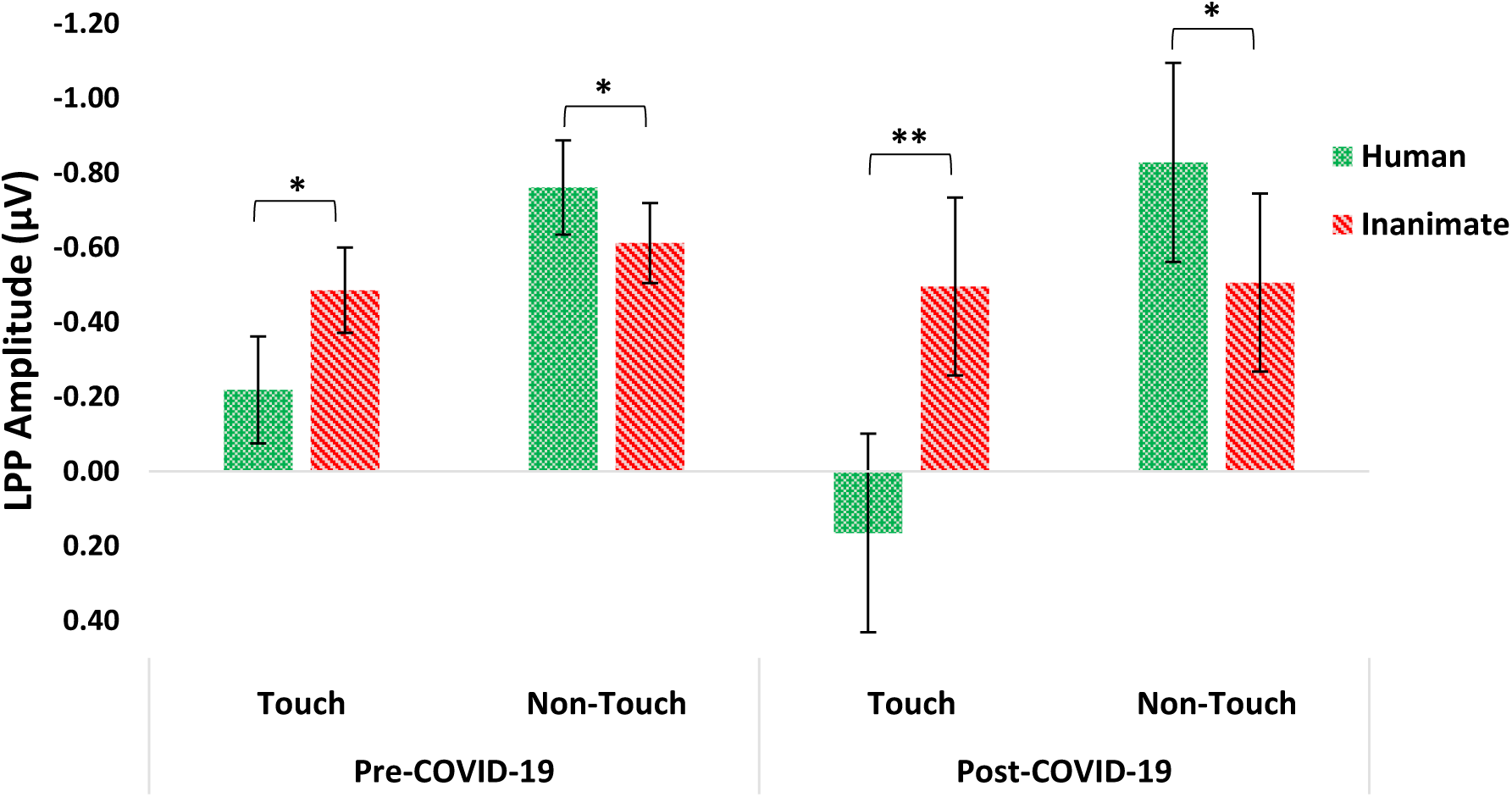
LPP average amplitudes for the pre- and post-COVID-19 groups (averaged between Fz, Cz, and Pz electrodes, 400–600ms post-stimulus). In both the pre- and post-COVID-19 groups, the LPP amplitudes were more positive during observation of human touch. The LPP amplitudes were also more positive when participants observed non-touch human photos than when they observed non-touch inanimate photos. A main effect was found for the touch condition. *Error bars represent standard error of the mean* \**P <.05* \*\**P*<.*001*

**Figure 7.**
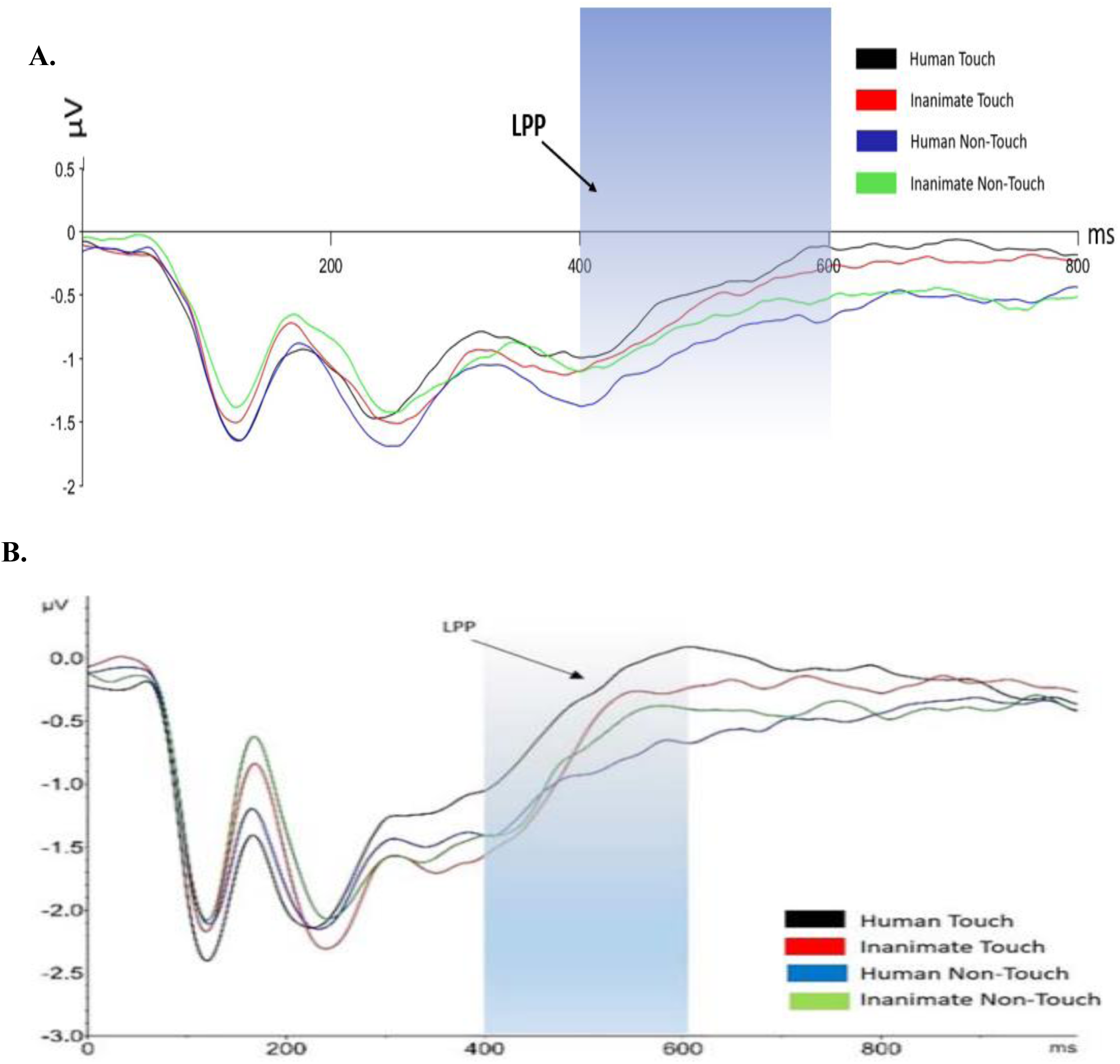
Onset-synchronized grand-average ERP LPP component measured under the indicated experimental conditions. The LPP component was averaged between the midline electrodes Fz, Cz, and Pz. The blue shaded areas indicate the time windows for LPP (400–600 ms post-stimulus). **(A)** LPP amplitudes for the post-COVID-19 group. **(B)** LPP amplitudes for the pre-COVID-19 group (adapted from Peled-Avron et al., 2016).

The findings also revealed a significant main effect for touch in both groups (pre- and post-COVID-19) (*F* (1,53) =26.481, *P*<.001, partial *η^2^*= .333; *F* (1,37) = 41.69, *P*<.001, partial *η^2^*= .53; respectively), indicating that human touch photos evoked higher LPP amplitudes than touch photos containing inanimate objects. Contrary to our hypothesis, the analysis did not reveal a significant interaction between touch and COVID-19 (*F* (1,90) = 2.74, *p* =.101, partial *η^2^* =.03) or between object and COVID-19 (*F* (1,90) =0.742, *p* =.391, partial *η^2^* =.008). On the other hand, we found a significant main effect for touch (*F* (1, 90) = 68.226 *P*<.001, partial *η^2^*=.431), such that the LPP amplitudes were more positive towards participants observed touch photos compared to non-touch photos. No significant main effect was found for object, (*F* (1, 90) = 3.182, *p* =.078, partial *η^2^*= .034).

## Discussion

In the current study, we examined how perceptions of interpersonal touch in social interactions were affected by the COVID-19 pandemic. Specifically, we compared pre- and post-COVID-19 behavioral and neural responses to observed social touch. The stimuli described in Peled-Avron et al. (2016; 2017) enabled us to compare behavioral and neural data (Peled-Avron et al., 2016; Peled-Avron & Shamay-Tsoory, 2017; respectively) collected prior to COVID-19 to a new set of data collected post-COVID-19. We hypothesized that after COVID-19 social touch might induce hypervigilance due to the risk of infection. Thus, we focused on the P1 and LPP ERP components, which respectively reflect early and late hypervigilance. As predicted, we found behavioral changes in social touch perception: participants in the post-COVID-19 group rated photos with *touch* as less pleasant than did participants in the pre-COVID-19 group. Moreover, participants in the post-COVID-19 group rated photos with *humans* as less pleasant than did participants in the pre-COVID-19 group. It is interesting to note that the post-COVID-19 group exhibited no differences in pleasantness ratings between human touch and inanimate touch photos, whereas for the non-touch condition, participants perceived photos of non-touching humans as more pleasant than photos of non-touching inanimate objects. This pattern was <u>not</u> found before COVID-19. In addition to behavioral changes, we found neural changes in the P1 amplitudes: In the pre-COVID-19 group, the P1 amplitudes were more positive when participants observed touch compared to non-touch photos. In contrast, the P1 amplitudes in the post-COVID-19 group did not exhibit any differences between touch and non-touch photos. Additionally, no differences were found in LPP amplitudes or in P1 latencies for the post-COVID-19 sample compared to the pre-COVID-19 sample.

The decreased pleasantness ratings for photos depicting touch and humans in the post-COVID-19 group may have been influenced by various factors associated with the pandemic. First, concerns about potential transmission of the virus through touch as well as increased awareness of hygiene practices may have led to the reduced pleasantness rating for touch-related stimuli. The changes in participants’ perceptions and evaluations of social touch and human stimuli after COVID-19 may represent a defensive response serving to mitigate potential risks and threats associated with the pandemic. By devaluing the pleasantness of touch and perceiving photos with human touch as less pleasant, individuals may be adopting a protective mechanism to avoid potential harm and reduce their susceptibility to the virus (Jones, 2007; Oaten et al., 2009). Moreover, these results may be aligned with the finding that perceived vulnerability to disease predicts conformist attitudes (Murray & Schaller, 2012). When collectively facing the threat of COVID-19, individuals may have adopted more cautious behaviors, conformed to public health guidelines and sought to minimize the risk of infection. Consequently, avoiding physical contact, including social touch, may be seen as a conformist response to the prevailing norms and expectations surrounding personal safety and health.

The results of the current study suggest that not only did the pandemic have an impact on behavioral responses, it also affected the neural processing of touch stimuli, thus generating a change in P1 amplitudes. Specifically, prior to COVID-19, stimuli depicting human touch elicited a stronger neural response than stimuli depicting humans who are not touching or stimuli depicting inanimate objects either touching or not touching. In contrast, after COVID-19 the differentiation between social touch and social non-touch diminished. These findings may reflect shifts in attention or changes in the salience of touch-related information due to the altered circumstances brought about by the pandemic. Moreover, these findings may be related to attentional bias. Attentional bias refers to the tendency to selectively attend to certain stimuli over others (Abado et al., 2020). In the context of touch perception and social stimuli, attentional biases can influence how individuals allocate their attention to these stimuli, subsequently affecting the P1 component (Clark & Hillyard, 1996). Prior to COVID-19, participants may have showed a bias in attention toward touch, as evidenced by the more positive P1 amplitudes for photos with touch compared to photos without touch. This may suggest that touch-related stimuli captured attention more effectively. Additionally, participants exhibited a bias toward photos depicting humans over those depicting inanimate objects, indicating a preference for attending to social stimuli. After COVID-19, however, the interaction between touch and object was not significant, indicating a reduction in attentional bias toward touch. This change may be attributed to altered attentional priorities or shifts in the salience of touch-related stimuli due to the COVID-19 pandemic. Overall, these findings suggest that attentional bias, as reflected in the early processing of visual stimuli (P1 amplitudes), may have been modulated by COVID-19 and its associated changes in perception and social interactions.

Another possible explanation for this shift may be associated with the notion that attention allocation leads to higher emotional processing (Li et al., 2014). Prior to the pandemic, when participants observed physical social touch they allocated more attention to the stimuli, as reflected in higher P1 amplitudes. This heightened attention, in turn, led to heightened emotional processing, as indicated by larger LPP values. After COVID-19, observing the same physical social touch did not evoke attention allocation (baseline P1 amplitudes) and this absence of attention did not lead to any change in LPP amplitudes (with LPP amplitudes still heightened in the presence of touch stimuli). Accordingly, a change in early attentional processing (reflected in P1 amplitudes) may be necessary for altering higher level emotional processing (reflected in LPP amplitudes). The observed shift in attention allocation may be related to the effects of the pandemic, which may have influenced participants’ cognitive and emotional responses during touch perception (see Figure 8).

**Figure 8.**
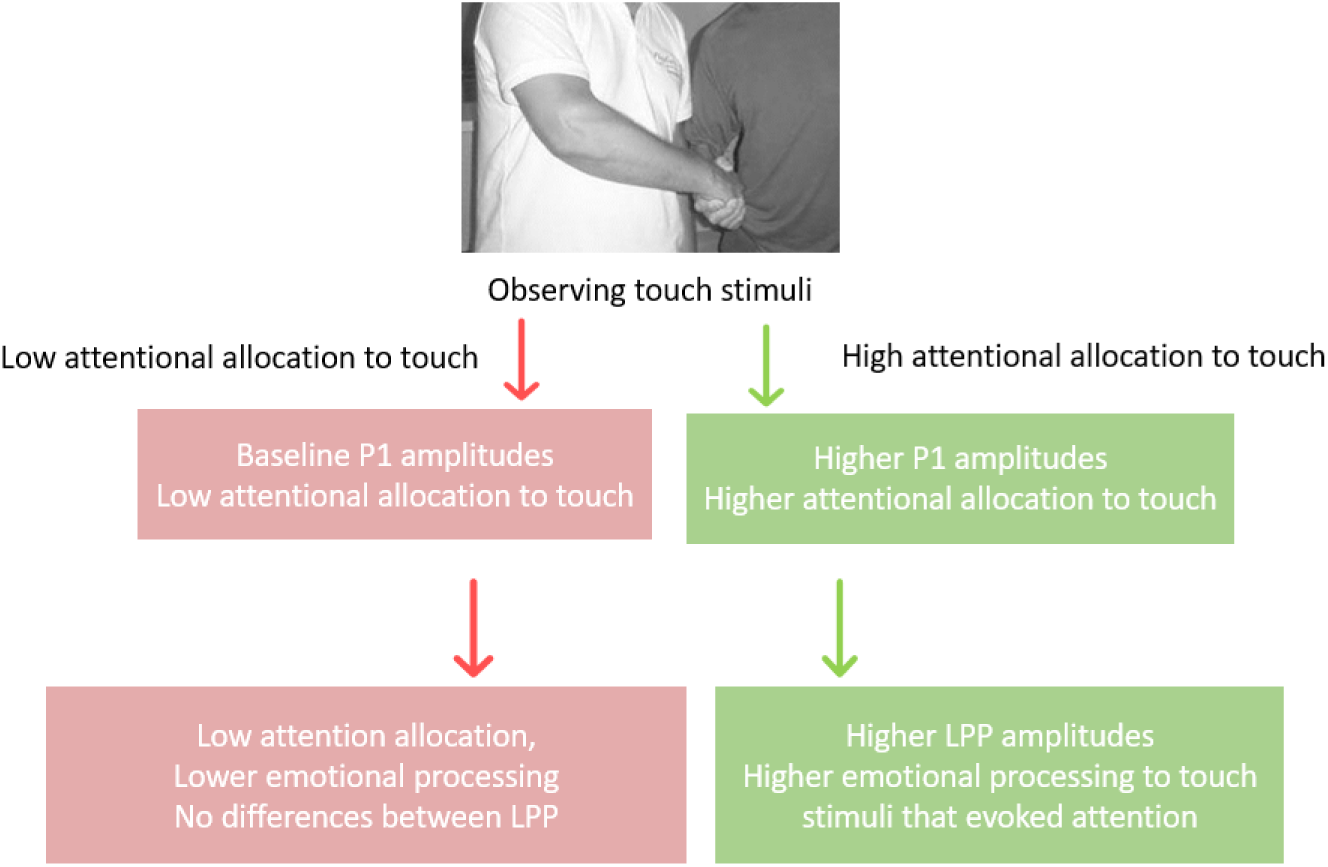
Possible explanation based on ERP results. When observing physical social touch, participants in the pre-COVID-19 group exhibited more attention allocation (larger P1 amplitudes), which in turn evoked heightened emotional processing (larger LPP amplitude). In contrast, for participants in the post-COVID-19 group, observing physical social touch did not evoke attention allocation (baseline P1 amplitudes). In the absence of a change in attention allocation, the LPP amplitudes did not change (still higher LPP in response to touch stimuli).

Moreover, Krusemark and Li (2013) showed that perception of disgusting pictures is associated with reduced P1 amplitudes, whereas neutral affective pictures elicit larger P1 amplitudes. This finding suggests that the P1 component is sensitive to emotional states, with neutral affective pictures increasing P1 amplitudes and disgusting pictures decreasing these amplitudes. In the context of our study, we can speculate that the observed decrease in P1 amplitudes when participants observed social touch photos may be attributed to feelings of disgust. The implication is that P1 amplitudes were less positive after COVID-19 than before the pandemic, possibly due to feelings of disgust when viewing the photos. This is noteworthy considering that disease prevention has been associated with the emotion of disgust (Jones, 2007; Oaten et al., 2009). To further enhance our understanding of the relationship between P1 amplitudes and perceived disgust in our study, it would have been beneficial to include a subjective rating task in which participants were asked to rate the photos on a scale specifically designed to measure the degree of disgust they experienced.

In the wake of the COVID-19 pandemic, it is important to examine the implications and impact of its social restrictions on human social functioning. Social touch is a crucial part of everyday social interactions. Whereas other researchers have examined perceptions of social touch during and after COVID-19, to our knowledge this study is the first to compare perceptions of social touch to a set of data collected prior to the pandemic, while also examining the neural mechanisms underlying these perceptions. The literature regarding the effects of COVID-19 on social touch is inconsistent. Some researchers claim that social restrictions may increase the craving for social touch in the same way that fasting increases the craving for food (e.g., Ujitoko et al., 2022, Von Mohr et al., 2021). Others argue that the social restrictions dictated by COVID-19 have changed how humans perceive touch, such that touch is now perceived as a threat signal that may even generalize into reduced positive reactions to observed touch (Massaccesi et al., 2021). Our results support the second argument and are in line with a study by Massaccesi et al. (2021), which demonstrated that images depicting crowds and large gatherings were rated as less positive during the pandemic period compared to the pre-pandemic period. Moreover, participants who reported overall greater physical isolation, stronger feelings of social closeness and higher perceived threat of COVID-19 gave more positive ratings to images depicting individuals alone and in very small groups. Our findings are also in line with Kühne et al. (2022), who found that following the pandemic, participants perceived situations in which two individuals were depicted wearing face masks and not interacting with one another as more positive. One important and surprising insight emerging from the current study is that COVID-19 has had an impact on human perception of social touch, such that observing nonsocial touch evoked more positive emotions than observing human touch.

Numerous studies have demonstrated that both observed touch and experienced touch activate similar brain regions (e.g., Ebisch et al., 2008; Keysers et al., 2004), suggesting that witnessing someone else being touched and experiencing touch oneself engage the same neural mechanisms (Schirmer et al., 2015). Nevertheless, it is crucial to recognize that the actual physical experience may differ, as observing photos may not fully capture the complex and dynamic nature of tactile interactions (Abraira & Ginty, 2013), especially those that occurred during the pandemic. The absence of physical and sensory experience in our study may limit the generalizability of our findings. Real-life encounters involving touch can evoke heightened feelings of threat or intimacy, potentially exerting an influence on individuals’ perceptions and responses that differs from the effect of static visual representations (Saarinen et al., 2021). Therefore, future research should consider incorporating real-life touch scenarios to gain a more comprehensive understanding and provide deeper insights into the complex interplay between social touch perception and COVID-19.

Overall, our results indicate that COVID-19 has modified how human beings perceive social touch. Our findings provide evidence for the impact of the pandemic on individuals’ perceptual and evaluative processes, highlighting the importance of considering social and environmental factors in understanding subjective experiences. Further research is needed to investigate the mechanisms underlying these effects and to determine the generalizability of these findings in different populations and contexts. Touch is vital for our social emotional development and provides us essential tools for expressing our emotions, establishing intimacy and maintaining social bonds (Harlow & Zimmerman, 1959). Therefore, studying and characterizing how the pandemic has altered social touch is crucial and can have a beneficial impact on how society functions as a whole.

